# Inner lumen proteins stabilize doublet microtubules in cilia/flagella

**DOI:** 10.1101/383430

**Authors:** Mikito Owa, Takayuki Uchihashi, Haruaki Yanagisawa, Takashi Yamano, Hiro Iguchi, Hideya Fukuzawa, Ken-ichi Wakabayashi, Toshio Ando, Masahide Kikkawa

## Abstract

Motile cilia are microtubule-based organelles that play important roles in most eukaryotes. Although it is known that microtubules in cilia are sufficiently stable to withstand their beating motion, it remains unknown how they are stabilized while serving as tracks for axonemal dynein and intraflagellar transport. To address this question, we identified a new class of microtubule-associated proteins, named FAP45 and FAP52, in *Chlamydomonas*. These proteins are conserved among eukaryotes with motile cilia. Using cryo-electron tomography (cryo-ET) and high-speed atomic force microscopy (HS-AFM), we established that lack of these proteins leads to a loss of inner protrusions in B-tubules and less stable microtubules. These inner protrusions are located near the inner junctions of doublet microtubules and lack of FAP45, FAP52, and FAP20 results in detachment of the B-tubule from the A-tubule, as well as flagellar shortening. These results demonstrated that FAP45 and FAP52 bind to the inside of microtubules and stabilize ciliary axonemes.

## Introduction

Cilia and flagella are microtubule-based organelles that operate as both antennae and propellers in eukaryotic cells. The structures and related genes of cilia and flagella are well conserved among eukaryotes. Cilia are classified as either non-motile or motile. Non-motile cilia, or “primary cilia”, function as antennae and are involved in signal transduction through the hedgehog, Wnt, and Ca^2+^ signaling^1^. Motile cilia and flagella beat at 20-60 Hz^2, 3^ and drive cellular motility and fluid flow. Since the cells in most tissues are ciliated, ciliary dysfunction leads to various types of human diseases termed ciliopathies, which include hydrocephalus, *situs inversus*, retinal degeneration, and nephronophthisis^4^. Furthermore, recent studies revealed that the assembly of cilia is significantly decreased in several types of tumors^5, 6^, suggesting a close relationship between cilia and tumorigenesis.

The axoneme is the core structure of cilia and flagella and is highly conserved from protists to vertebrates. The axoneme is composed of a central pair of microtubules cylindrically surrounded by nine doublet microtubules (DMTs). This arrangement is often referred to as the 9+2 structure (Fig. 1a). In DMTs, 10 protofilaments of the B-tubule are attached to 13 protofilaments of the A-tubule at the inner and outer junction (Fig. 1a). In motile cilia and flagella, structures essential for motility, such as axonemal dyneins, radial spokes, and the nexin-dynein regulatory complex (N-DRC), are arranged on DMTs with a 96-nm repeating unit^7-9^. Axonemal dyneins bound on A-tubules slide on the neighboring B-tubules and this sliding propagates along the axoneme, generating a bending force. The activity of dyneins is regulated by the interaction between the radial spokes and the central pair of singlet MTs^10-12^.

**Figure 1.**
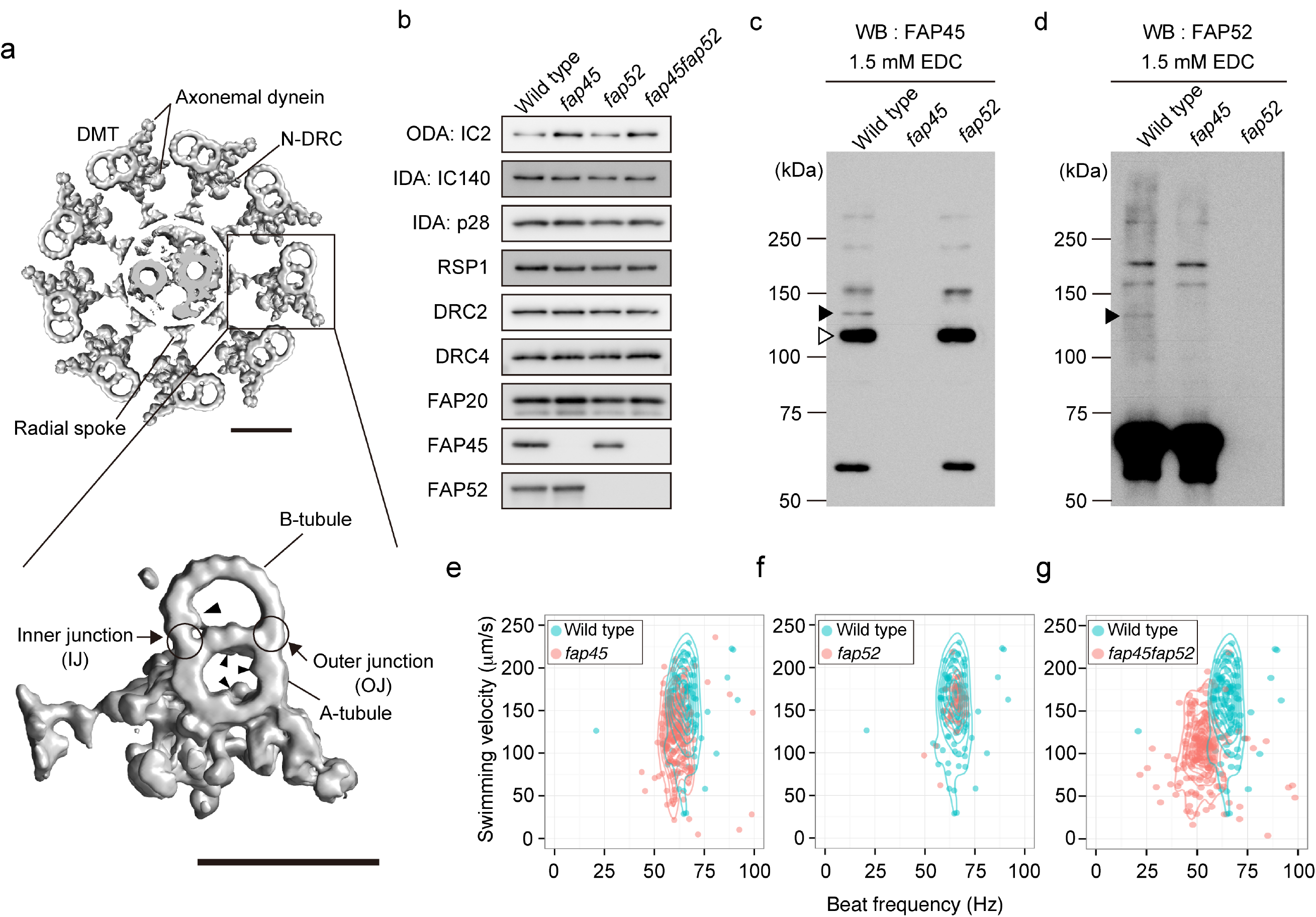
Characteristics of the FAP45 and FAP52 null mutants. (a) Top: a transverse view of the 9+2 structure of the *Chlamydomonas* axoneme. Scale bar = 50 nm. Bottom: a magnified DMT. Arrowheads indicate MIPs. Scale bar = 50 nm. (b) Western blot analyses of wild type, *fap45*, *fap52*, and *fap45fap52* double mutant axonemes stained with various antibodies. FAP45 and FAP52 proteins were not detected in *fap45* and *fap52*, respectively. Proteins essential for flagellar motility (ODA-IC2: outer dynein arm-intermediate chain 2; IDA-IC140: inner dynein arm-intermediate chain 140; IDA-p28: inner dynein arm-light chain p28; RSP1: radial spoke protein 1; DRC2 and 4: dynein regulatory complex 2 and 4; FAP20: inner junction protein of DMT) were not reduced in the mutants. (c, d) Axonemes from wild type and mutant *Chlamydomonas* crosslinked using EDC (zero-length crosslinker) were immunoblotted with anti-FAP45 (c) and anti-FAP52 antibodies (d). Filled arrowheads indicate the crosslinked product of FAP45 and FAP52 in a 1:1 ratio. The open arrowhead indicates the crosslinked product of FAP45 and tubulin. (e-g) Motility phenotypes of the mutants were assayed using the CLONA system. Swimming velocity and beat frequency were slightly reduced in *fap45*, whereas no significant reduction was observed in *fap52*. *fap45fap52* showed a more severe phenotype than did *fap45*.

Recent advances in cryo-ET techniques have dramatically revealed structural details of DMTs, yet a significant question remains—What stabilizes DMTs? Cytoplasmic MTs frequently bend with various degrees of curvature^13, 14^ and this bending occasionally induces MT breakage^15^. Furthermore, a recent study demonstrated that cytoplasmic MTs were damaged even after bending only several times^16^. Therefore, to ensure a high beat frequency, the DMTs of motile cilia and flagella should be structurally robust compared with cytoplasmic MTs. However, the mechanism that provides DMTs with such robustness remains to be clarified.

Nicastro and colleagues have provided insights into the structural basis of DMTs using cryo-ET of *Chlamydomonas* flagella. They reported periodic high densities on the inner surfaces of A-tubules and B-tubules, which they named microtubule inner proteins (MIPs, Fig.1a) ^7, 17^. MIPs have also been observed in the axonemes of higher organisms ^7, 17-20^, implying that these inner structures are essential for the integrity of DMTs in motile cilia and flagella.

In this paper, we explored the mechanisms stabilizing DMTs using *Chlamydomona*s mutants and various techniques, including cryo-ET and HS-AFM. We found that two uncharacterized flagellar-associated proteins (FAP), FAP45 and FAP52, are essential for the stability of B-tubules. Lack of both FAP45 and FAP52 leads to the loss of MIPs in B-tubules, resulting in disruption of the MT walls. Furthermore, the B-tubule wall detached from the A-tubule at the inner junction, and flagellar shortening was observed when both FAP52 and FAP20 were absent. These results indicate that the B-tubule is reinforced by microtubule inner proteins and its stability is essential for maintaining the structure of the axoneme.

## Results

### Identification of proteins essential for DMT stabilization in flagella

To identify proteins that stabilize DMTs, we searched the *Chlamydomonas* flagellar proteome database^21^ using the assumption that these proteins are (1) abundant in the axoneme fraction, (2) tightly bound on the axoneme even in the presence of high salt, and (3) highly conserved among ciliated organisms. The proteins that satisfy these criteria are listed in Supplementary Table S1. We focused on FAP45, an uncharacterized protein whose peptides were most frequently found in the proteomic analysis (Table S1, “total unique peptide” and “Axo” columns). FAP45 is a coiled-coil protein composed of 501 amino acids, has a predicted molecular mass of a ~59 kDa, and is conserved among organisms with motile cilia. The human ortholog of FAP45 is coiled-coil domain-containing protein 19 (CCDC19), also known as NESG1. The transcript of this protein is enriched in the nasopharyngeal epithelium and trachea^22^. However, the functions of FAP45/CCDC19 are totally unclear and thus we focused on FAP45.

We first investigated the partner that interacts with FAP45 on the axoneme using chemical crosslinking. We treated isolated wild type axonemes with 1-ethyl-3-(3-dimethylaminopropyl)carbodiimide (EDC, zero-length crosslinker). The crosslinked products were solubilized and immunoprecipitated with a polyclonal anti-FAP45 antibody. Tubulin was identified as the major crosslinked partner of FAP45 using western blot (Fig. 1c, open arrowhead; data not shown), suggesting a direct interaction between FAP45 and tubulin. In addition, a mass spectrometric analysis revealed that the ~130 kDa product is composed of FAP45 and FAP52 proteins, probably in a 1:1 ratio (Fig. 1c and d, filled arrowhead; Supplementary Table S2).

FAP52 is the *Chlamydomonas* ortholog of human WD40 repeat domain 16 *(WDR16)* and is listed in Supplementary Table S1. It was previously shown that *WDR16* knockdown in zebrafish led to severe hydrocephalus^23^. Furthermore, a recent study reported that homozygous deletion of the *WDR16* gene in human leads to situs anomalies, which are typical phenotypes of ciliopathy^24^. However, the function of the FAP52/WDR16 protein in flagellar motility and axonemal structure remains unclear. Therefore, we hypothesized that FAP45 and FAP52 proteins play important roles in stabilizing DMTs.

### Isolation and characterization of *Chlamydomonas* FAP45 and FAP52 mutants

We used *Chlamydomonas* mutants lacking FAP45 and FAP52 to investigate their function. The mutants were isolated from ~10,000 clones in an insertionally-mutagenized *Chlamydomonas* library^25^. In these mutant strains, the ~1.8 kbp *aphVII* fragment was inserted into the 5’UTR of the FAP45 gene or the first exon of the FAP52 gene (Supplementary Fig. S1b). These strains did not express detectable FAP45 or FAP52 protein as analyzed by a western blot of axonemal fractions (Fig. 1b, Supplementary Fig. S1c). On the other hand, *fap45* and *fap52* axonemes retained the known major axonemal components, such as outer arm dyneins, inner arm dyneins, radial spokes, and the dynein regulatory complex (N-DRC) (Fig. 1b). Despite FAP45 and FAP52 being crosslinked by a zero-length crosslinker, they localized on the axoneme independent of each other (Fig. 1b). Consistent with the mass spectroscopic analysis, the ~130 kDa crosslinked product of FAP45 and FAP52 was not detected in the crosslinked *fap45* or *fap52* axoneme (Fig. 1c and d, filled arrowheads), suggesting that FAP45 and FAP52 are neighbors on the axoneme.

Next, we examined the motility phenotype of *fap45* and *fap52*. The swimming velocity and beat frequency of *fap45* cells were slightly reduced (Fig. 1e, Movie 1), whereas the phenotype of *fap52* was quite similar to that of wild type (Fig. 1f, Movie 1). Intriguingly, the double mutant *fap45* and *fap52 (fap45fap52)* swam significantly slower than wild type, with a low beat frequency (Fig. 1g, Movie 1), even though there was no decrease in the major known axonemal components essential for flagellar motility (Fig. 1b). Furthermore, there was an accumulation of non-motile *fap45fap52* cells under confluent culture conditions (Movie 2). Since the medium under confluent culture conditions is more viscous than that of log phase conditions, a viscous load may affect the motility of *fap45fap52*. To explore this possibility, we observed the swimming behavior of wild type and *fap45fap52* log phase cells in this viscous medium. Most of the wild type cells swam slowly but smoothly (Movie 3) whereas many of the *fap45fap52* cells stopped swimming or struggled to swim against the viscous load (Movie 3), indicating that *fap45fap52* flagella cannot beat properly under these conditions. Since the *fap45fap52* axonemes had normal levels of axonemal dyneins, radial spokes, and N-DRC, these results suggest that the lack of both FAP45 and FAP52 causes structural defects in the DMT.

### FAP45 and FAP52 are luminal proteins in B-tubules

We biochemically localized FAP45 and FAP52 in DMTs by fractionating the axonemal proteins using increasing concentrations of sarkosyl^26^ (Supplementary Fig. S2a). FAP45 began to be extracted from the pellet at 0.2% sarkosyl, half of the FAP45 was extracted at 0.3%, and the protein was completely extracted from the pellet at 0.7% (Supplementary Fig. S2b). FAP52 started to be solubilized at 0.3% sarkosyl and almost all the protein was solubilized at 0.7%, although a small amount of protein still remained in the pellet. These behaviors correlate well with solubilization of the B-tubule.

To distinguish whether FAP45 and FAP52 are located outside or inside the B-tubule, we prepared axonemes containing biotinylated FAPs and tested whether or not the proteins are accessible to streptavidin. These axonemes were purified from rescue strains expressing FAP proteins whose N- or C-terminus was fused to biotin carboxyl carrier protein tag (BCCP tag, Fig. S2b and c) ^27, 28^. However, essentially no signal of biotinylation was detected on those axonemes (Supplementary Fig. S2d), suggesting that streptavidin access to BCCP tag was prevented, presumably by the B-tubule wall. Therefore, we treated axonemes with 0.15% sarkosyl, which partially broke the B-tubule wall and, as expected, resulted in detection of the tagged proteins (Supplementary Fig. S2e). Taken together with the results that the loss of FAP45 and FAP52 did not affect the incorporation of other axonemal proteins bound on the outer surface of the DMT (Fig. 1b), these data indicate that FAP45 and FAP52 are luminal proteins enclosed by the B-tubule.

### Lack of FAP45 and/or FAP52 causes structural defects in B-tubules

To reveal the defect caused by the loss of FAP45 and FAP52, we observed the mutant axonemes by cryo-ET. Interestingly, the mutants had structural defects inside of the B-tubule (Fig. 2). For further visualization of missing structures in *fap45* and *fap52*, we also applied Student’s *t*-tests to compare wild type and mutant density maps^27^ (Fig. 2e and f, right).

**Figure 2.**
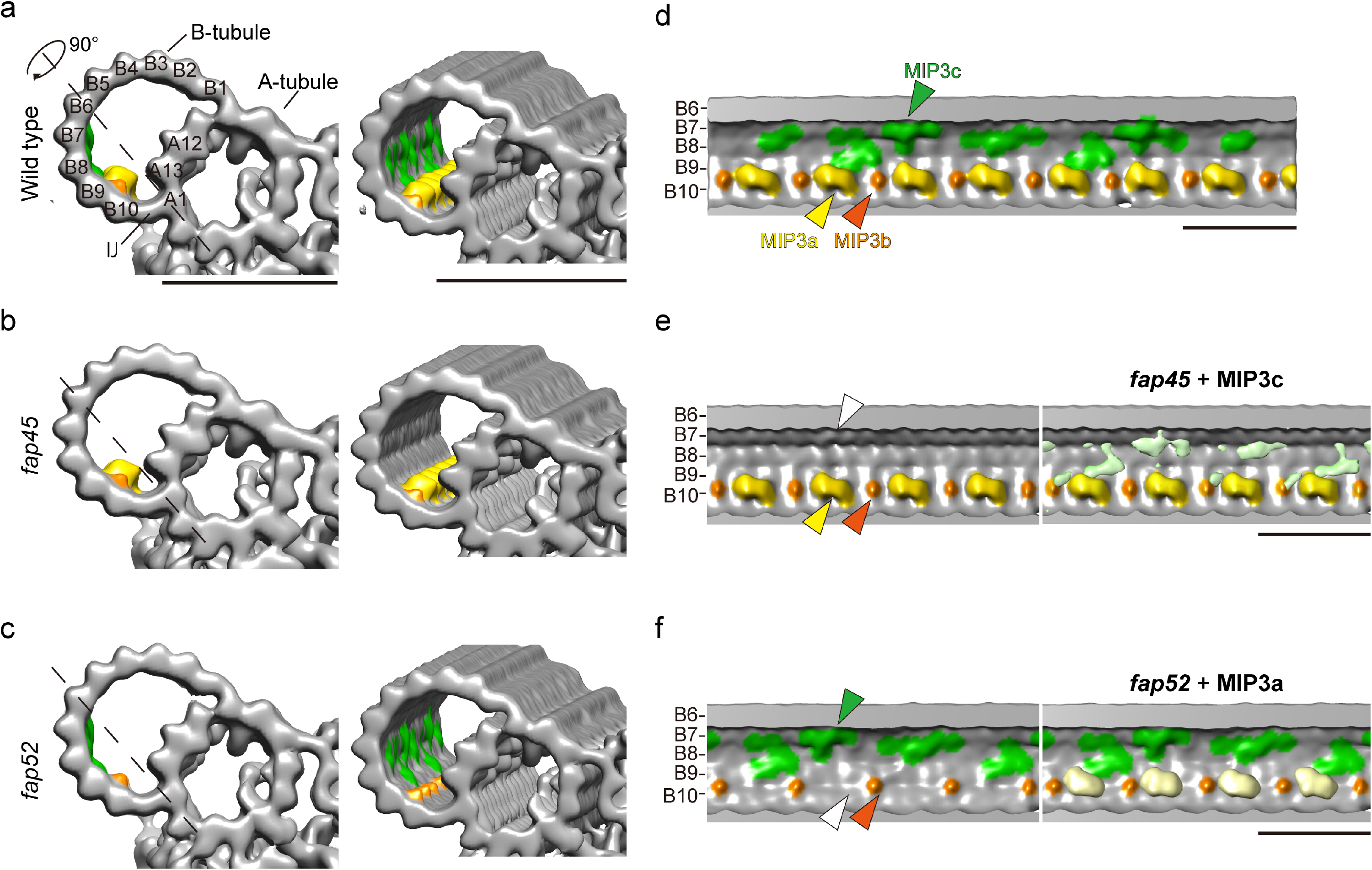
Cryo-ET showed that axonemes of *fap45* and *fap52* mutant have structural defects in B-tubules. Isosurface renderings of averaged axonemal 96-nm repeats from wild type (a, d), *fap45* (b, e), and *fap52* (c, f). a-c are cross-sectional and oblique, and d-f are longitudinal views. The black dashed lines in a-c indicate the slicing plane of d-f. Protofilament numbers are labeled in a and d-f. Missing structures in *fap45* and *fap52* were determined by Student’s t-tests as previously described^27^. The t-value maps acquired by the t-tests were overlaid on the renderings of *fap45* (e, light green) and *fap52* (f, light yellow). In this figure, MIP3a and b were colored by yellow and orange, respectively. MIP3c (the missing structure in *fap45*) was colored green, according to the *t*-value map shown in e. Scale bar = 25 nm.

A comparison between wild type and *fap45* DMTs shows that the luminal surface of wild type B-tubule is smoother than that of *fap45* (Fig. 2d and e). In the *fap45* DMTs, the grooves between B-tubule protofilaments B7-B9 are prominent (Fig. 2b and e; Supplementary Fig. S3e), which are similar to bare microtubules. The grooves seem to be partially filled with filamentous densities as shown in the t-map (Fig. 2e right, highlighted in green). These densities are arranged in a 48-nm repeat, and reminiscent of filamentous densities extended from MIP3 in *Tetrahymena* DMTs^29^. In both our *Chlamydomonas* DMT and *Tetrahymena* DMT^28^, these lateral filaments appear to connect MIP3 to protofilaments B7-B9. Therefore, we named these densities that are missing in *fap45* mutant as “MIP3c” (Fig 2e right and Fig. 6a).

**Figure 6.**
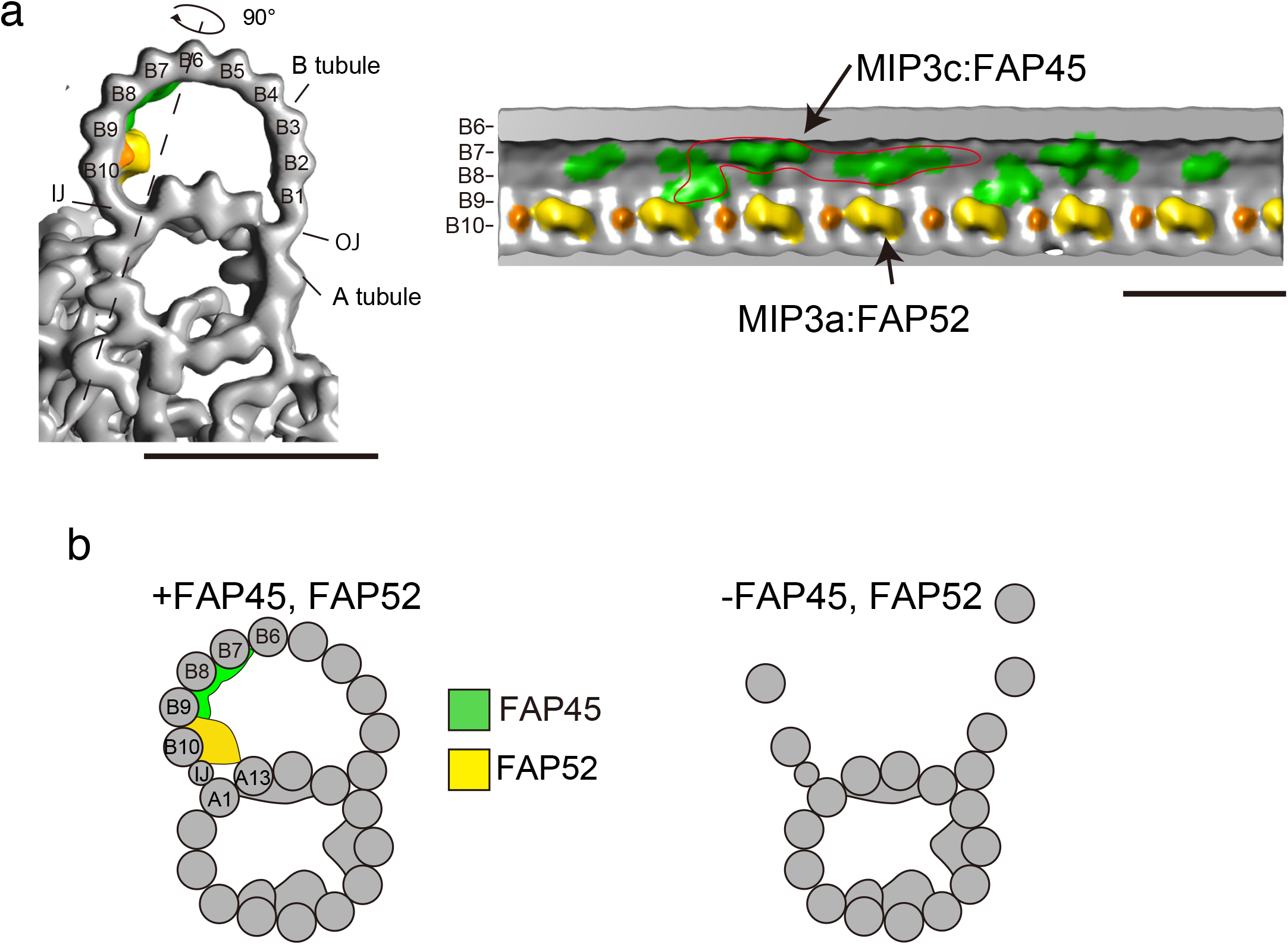
A schematic model of MIP3. (a) Isosurface rendering of the wild type density map. MIP3a-c is highlighted as shown in Fig. 2 (MIP3a: yellow; MIP3b: orange; MIP3c: green). The right map is the longitudinal view sliced by the line in the left map. A possible unit of MIP3c is circled in red. Scale bar = 25 nm. (b) A model of DMT stabilization. FAP45 (green) and FAP52 (yellow) are localized on the inner lumen of the B-tubule and stabilize the MT wall. In particular, FAP45 ties protofilaments B6-B9. FAP52 anchors B9 and B10 to A13 and binds to FAP45. Thus, these proteins strengthen the MT wall against DMT-bending. Loss of both proteins leads to destabilization of the B-tubule and the B-tubule is depolymerized by bending.

A comparison between the wild type and *fap52* DMT revealed an arch-like density is missing in the B-tubule of the *fap52* mutant. In the wild type DMT, there are a large arch-like density and a small spike-shaped density in a 16-nm repeat inside of the B-tubule on the protofilament B9 and B10, called MIP3a and MIP3b, respectively (Fig. 2a and d, highlighted in yellow and orange; Supplementary Fig. S3d)^17^. The DMTs of *fap52* completely lacked the densities of MIP3a, whereas MIP3b appear to be unchanged (Fig. 2c and f; Supplementary Fig. S3d).

Next, we tested whether the predicted atomic structures of FAP45 and FAP52 can fit into these densities. *Chlamydomonas* FAP52 contains eleven WD40 domains (Supplementary Fig. S1a). Since WD40 repeat domains typically form a seven-bladed β-propeller, FAP52 most likely contains two β-propellers. Indeed, a model comprising two β-propellers fit well into the density map, suggesting that MIP3a corresponds to most of FAP52 protein (Supplementary Fig. S3g). For MIP3c, predicted coiled-coils of FAP45 protein were fit into the density (Supplementary Fig. S3f). FAP45 is mainly composed of helices and coiled-coils that account for 85% of the total sequence (Supplementary Fig. S1a). The total length of helices and coiled coils in FAP45 is ~60 nm. Therefore, the circled densities in Supplementary Fig. S3f are probable one unit of MIP3c. Thus, MIP3a and MIP3c most likely consist of FAP52 and FAP45, respectively.

To investigate the reasons why the double mutant *fap45fap52* cells swim significantly slower than single mutants, we observed the *fap45fap52* axoneme using cryo-ET. In this case, we could not apply 3D sub-tomographic averaging because many of the axonemes were frayed and the B-tubules were partially missing (Fig. 3a and b). Since cryo-ET is not suitable for observing the shapes of individual DMTs due to the missing wedge effects, thin sections of the *fap45fap52* axoneme were observed by conventional EM and revealed that the B-tubules were missing in ~33.8% of the outer DMTs (Fig. 3c). All of the protofilaments in the B-tubules were completely missing in some DMTs, whereas DMTs remaining several protofilaments in the B-tubules were also observed. These data suggest that the loss of both FAP45 and FAP52 decreases structural stability between the B-tubule protofilaments. Of note, we also found that the density of dynein e, a species of inner arm dyneins was decreased in *fap45* and *fap52* (Supplementary Fig. S3a-c). The density of dynein e is smaller than that of other inner arm dyneins even in wild type (Supplementary Fig. S3a), implying that binding of dynein e to the B-tubule is flexible compared to that of other dyneins. Therefore, it is possible that the lack of FAP45 or FAP52 slightly changes the shape of B-tubules and affects the interaction between dynein e and B-tubules.

**Figure 3.**
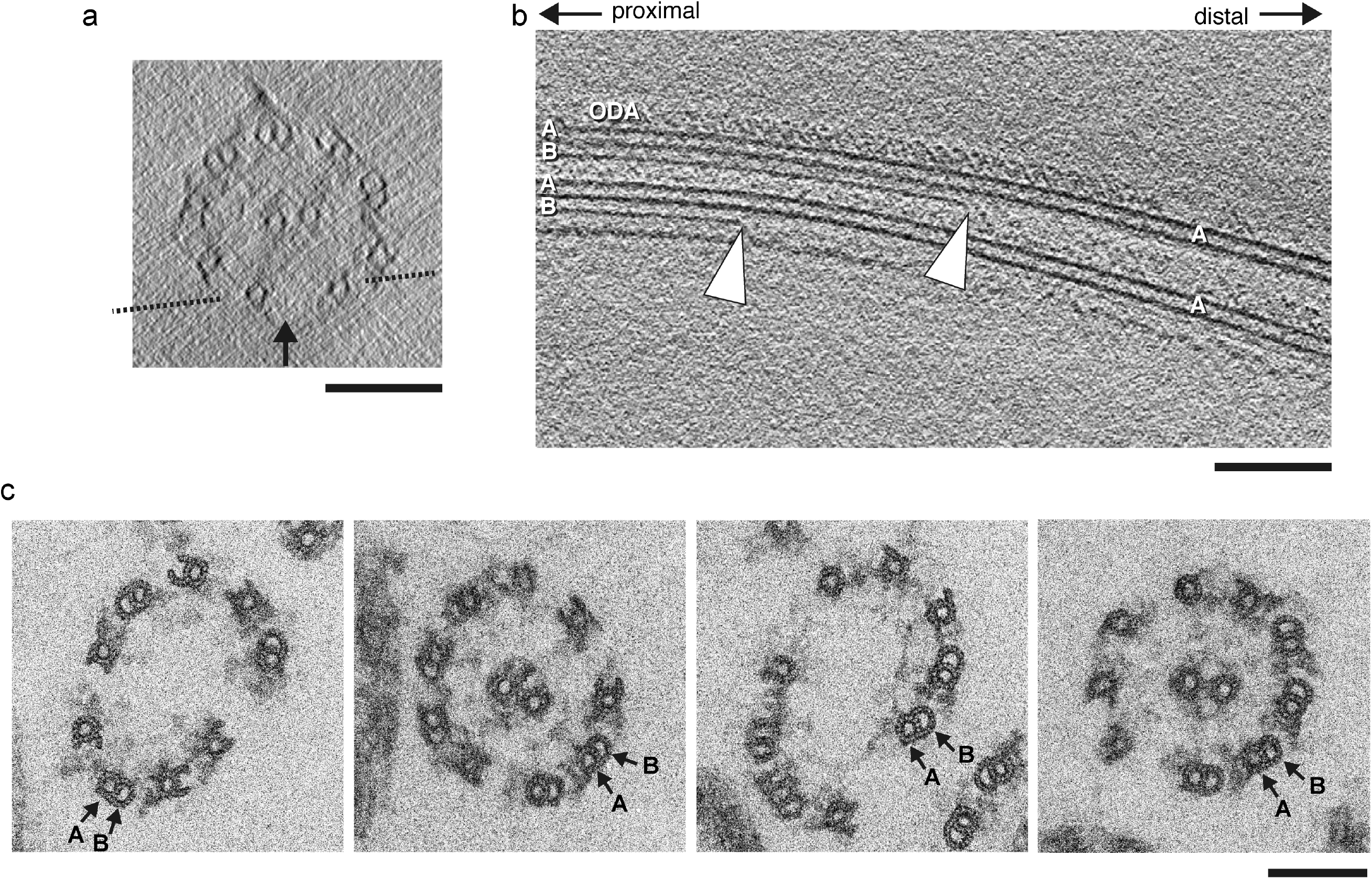
B-tubules are partially depolymerized in *fap45fap52*. (a, b) A typical tomogram of a *fap45fap52* axoneme in cross-sectional (a) and longitudinal view (b). A dashed line in a indicates the slicing plane of b. Arrowheads in b show break points in the B-tubules. (c) Typical thin section TEM images of *fap45fap52* axonemes. Broken B-tubules were observed in 33.8% of the DMTs (n = 552). The 9+2 structure was disorganized and the axonemes were frayed in regions containing broken B-tubules. Scale bar = 100 nm.

### Direct observation of B-tubule depolymerization by HS-AFM

The above “static” DMT structures suggested that the B-tubules of *fap45fap52* are less stable than those of wild type. We directly observed the stability of the B-tubule using HS-AFM^30^, in which a small cantilever tip (tip diameter ~ 1-2 nm) intermittently taps the DMT surface fixed on a mica stage. This intermittent contact can eliminate frictional force during lateral scanning and damage to the DMT caused by scanning because the feedback operation maintains a constant tip-sample interaction force and the force is applied for a short time (< 100 ns).

We first observed in vitro polymerized MTs in the absence of a stabilizing agent such as taxol. It was previously reported^25^ that MTs depolymerize spontaneously within two or three frames (1.0 to 1.5 s), suggesting that depolymerization requires less than 50 nm/s. Actual depolymerization could be faster^26^ but our HS-AFM has insufficient temporal resolution to verify this.

Compared with in vitro polymerized MTs, DMTs are far more stable and do not depolymerize spontaneously under normal HS-AFM conditions. To determine how a defect influences the stability of DMTs, we made a small hole by thrusting the tip of the cantilever into the B-tubule (Time=0, Fig. 4a) and observing whether or not the damage propagates. In wild type, the hole rarely grew during the observation period (Fig. 4b, Movie 4). On the other hand, in the mutants, the hole gradually expanded in both the proximal and distal directions (Fig. 4c, Movie 5-7). Time-course analysis of the depolymerized B-tubule length demonstrated that the tendency for depolymerization followed the order as *fap45fap52* > *fap45* > *fap52* ≥ wild type (Fig. 4d). These data strongly suggest that FAP45 and FAP52 stabilize the doublet B-tubule and prevent depolymerization induced by damage to the MT wall.

**Figure 4.**
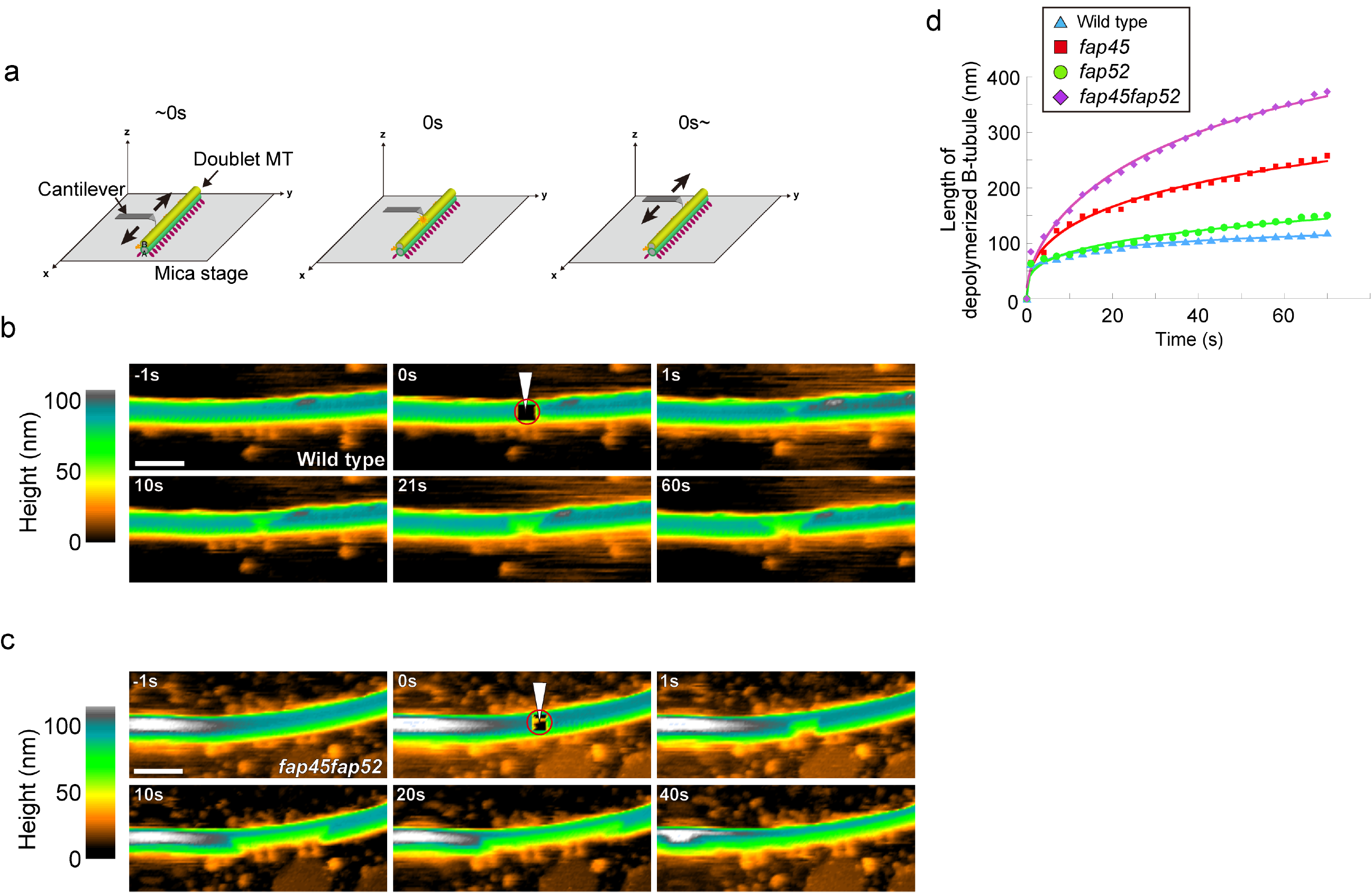
Loss of FAP45 and FAP52 results in destabilization of B-tubules, as observed by high speed-AFM. (a) Schematic of high speed-AFM observation of the DMT. The tip of the cantilever (tip diameter ~1-2 nm) intermittently tapped the DMT surface on a mica stage. At time 0s, the tip of the cantilever was thrust into the B-tubule and made a small hole (circled in red in b and c), immediately followed by normal HS-AFM observation. (b, c) Representative time-lapse images of wild type (b) and *fap45fap52* (c) DMTs acquired by HS-AFM. The images are colored according to the height from the mica stage. In the wild type, enlargement of the hole usually stopped within a few seconds, whereas most of the B-tubules in the field were broken in *fap45fap52* after 40s. Scale bar = 100 nm. (d) The average length of the depolymerized B-tubules was plotted against time (wild type: n = 11, total length = 7.25 μm; *fap45:* n = 8, total length = 4.92 μm; *fap52:* n = 7, total length = 3.83 μm; *fap45fap52*: n = 10, total length = 4.95 μm).

### MIP3a is important for B-tubule anchoring to the A-tubule at the inner junction of DMT

Since MIP3a is located near the inner junction between the A- and B-tubules and appeared to attach to the A-tubule^28^, we investigated whether FAP52 is involved in stabilizing the junction. We previously reported that FAP20 is a component of the inner junction and a null mutant of FAP20 *(fap20)* showed an abnormal swimming phenotype, although the structure of the DMTs appeared to be normal^31^. Based on the hypothesis that MIP3a anchors the B-tubule to the A-tubule together with FAP20, we constructed the double and triple mutants *fap20fap45*, *fap20fap52*, and *fap20fap45fap52*. Most *fap20fap45* cells were trembling and their motility appeared slightly worse than that of *fap20* cells (Movie 8). In contrast, *fap20fap52* and *fap20fap45fap52* cells were essentially paralyzed (Movie 8). Furthermore, the flagellar lengths of *fap20fap52* and *fap20fap45fap52* deviated greatly from wild type, *fap20*, and *fap20fap45* (Fig. 5c), with a far higher ratio of short flagella. To identify the cause of these defects, we observed the axonemes of the mutants by thin section TEM. The axonemes and DMTs of *fap20fap45* appeared similar to those of *fap20* (Fig. 5c and d). Interestingly, several doublet B-tubules in *fap20fap52* and *fap20fap45fap52* were detached from the A-tubules (Fig. 5c arrowheads; Fig. 5d) whereas no detachment of B-tubules from A-tubules was observed in *fap20fap45* axonemes. Given that such defects were rarely observed in *fap20* axonemes^31^, these results demonstrate that MIP3a is required for anchoring the B-tubule to the A-tubule at the inner junction together with FAP20, and the junction is important for stabilizing the axonemal structure.

**Figure 5.**
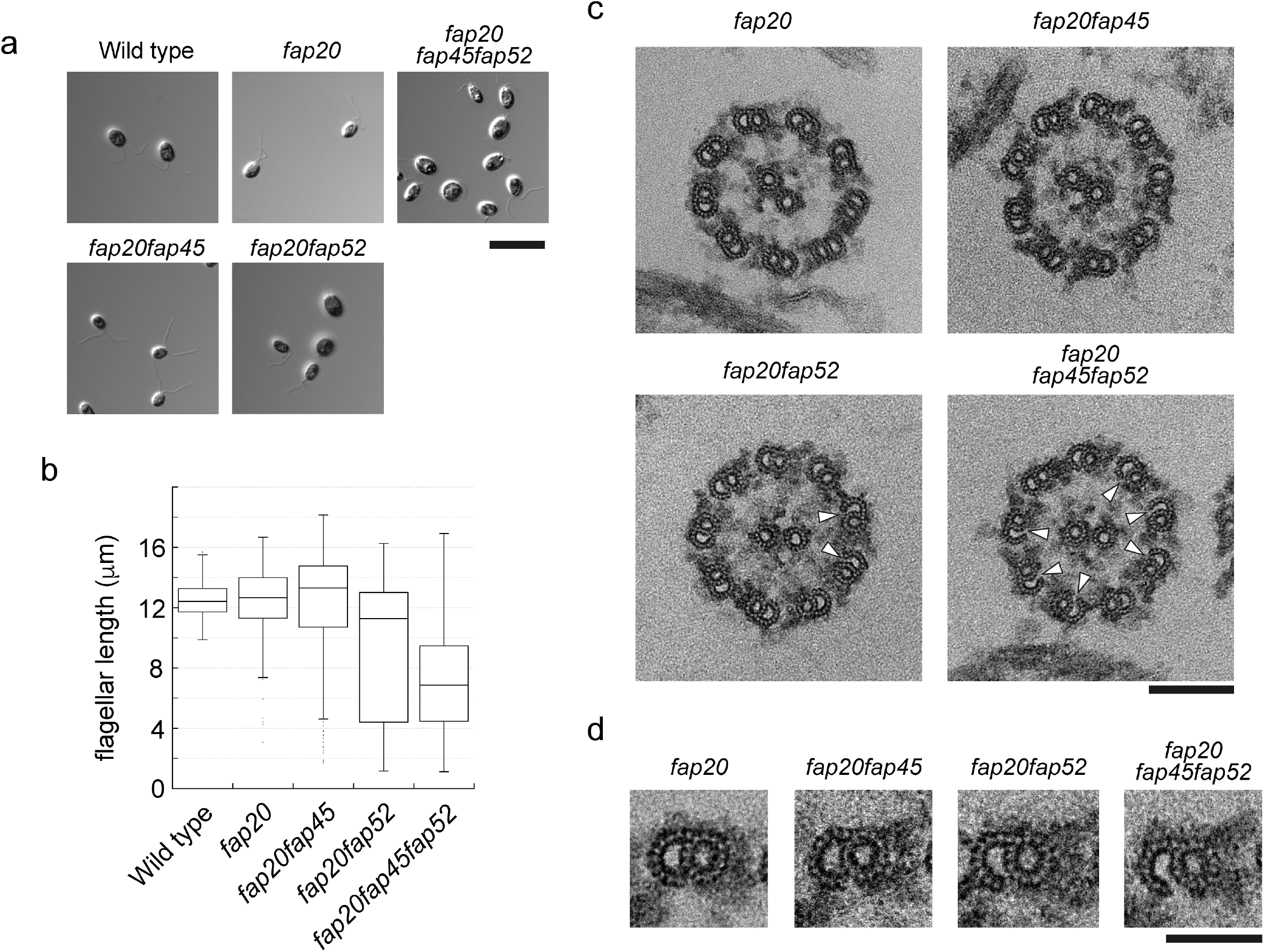
MIP3 is important for B-tubule anchoring to the A-tubule at inner junctions. (a) Differential interference contrast (DIC) images of wild type, *fap20*, *fap20fap45*, *fap20fap52*, and *fap20fap45fap52* cells. Scale bar = 10 μm. (b) Box plots of flagellar length in wild type, *fap20*, *fap20fap45*, and *fap20fap52*. *fap20fap52 and fap20fap45fap52* flagella were significantly shorter than in wild type, fap20, and fap20fap45 (Scheffe’s test). (c) Cross-sectional EM images of *fap20, fap20fap45, fap20fap52*, and *fap20fap45fap52* axonemes. Arrowheads indicate disconnections between the A- and B-tubules. Scale bar = 100 nm. (d) Cross sectional images of *fap20*, *fap20fap45*, *fap20fap52*, and *fap20fap45fap52*DMTs. Scale bar = 50 nm.

## Discussion

Ciliary/flagellar MTs are remarkably stable compared with cytoplasmic MTs. Of the “9+2” MTs, DMTs are directly stressed by the axonemal dynein-generated force and constantly bent and straightened in *Chlamydomonas* flagella or trachea cilia, yet their structures remain intact. The mechanism that stabilizes axonemal MTs has not been clarified. Here, we identified FAP45 and FAP52 as proteins that stabilize DMTs by binding to the inner lumen of the B-tubule. This is the first report to identify factors essential for the stabilization of ciliary/flagellar MTs.

Based on our studies and existing data, we propose a schematic model of FAP45 and FAP52 (Fig. 6b). Our cryo-ET observations show that MIP3a and MIP3c are composed of FAP52 and FAP45, respectively (Fig. 6a). FAP52/MIP3a anchors the protofilaments A13 and B10, whereas FAP45/MIP3c is bound inside of B7, B8, and B9 protofilaments. Besides, biochemical crosslinking shows that FAP45 and FAP52 directly interact with each other. Thus, these two proteins bundle B7-B10 protofilaments to reinforce the B-tubule. Consistent with this model, the B-tubules of *fap45* were more vulnerable to physical stress than those of *fap52* in the HS-AFM observation (Fig. 4d, Movie 6). The MIP3a structure was observed also in vertebrates^7, 18^, but MIP3c was not clearly described probably due to the limitation of resolution. However, the amino acid sequence of FAP45 is highly conserved from protists to mammals (Supplementary Fig. S4), suggesting that the functions of FAP45 are conserved also in higher organisms as in *Chlamydomonas*.

Our data also suggest that DMTs are stabilized by “fail-safe” mechanisms. The single mutants of FAP45 and FAP52 did not show significant decreases in swimming speed and only the double mutant *fap45fap52* showed slow swimming and defects of the B-tubules. Although the B-tubules of *fap45fap52* were more easily depolymerized than wild type B-tubules, the speed of depolymerization was slower than that of in vitro polymerized pure MTs. This suggests that B-tubules are also stabilized by other mechanisms, such as tubulin acetylation^32^ (Supplementary Fig. S5) or fMIPs that bind inside the B-tubule along its length^29^.

The inner junctions of DMTs are also stabilized by “fail-safe” mechanisms involving multiple proteins. We previously demonstrated that FAP20 is a constituent of the inner juction^31^. In addition to the “true” inner junction, our data clarified that MIP3a also connects B-tubules to A-tubules (Fig. 5c and d). Furthermore, some inner junctions appeared to be attached in *fap20fap52* and *fap20fap45fap52* mutants (Fig. 5c). In the triple mutant, additional proteins, such as tektin and PACRG, may contribute to stabilizing the inner junction, given our previous observation that these two proteins were partially retained in *fap20*^31^.

Ciliopathy caused by FAP45/CCDC19 mutation has not been reported to date. On the other hand, patients lacking FAP52/WDR16 were reported to have abdominal situs inversus and situs inversus totalis^24^. Despite abnormal laterality, those patients did not have typical symptoms of primary ciliary dyskinesia, such as recurrent bronchiolitis, hydrocephalus, or low nasal nitric oxide levels. These findings are consistent with the *fap52* phenotype in *Chlamydomonas*, where we could not detect abnormal phenotypes other than the lack of MIP3a structure. Given that the phenotype of *fap45* was more severe than that of *fap52* in *Chlamydomonas*, we predict that the lack of FAP45/CCDC19 in human causes ciliopathy.

Compared to B-tubules, the functions of MIPs in the A-tubules are unclear. A recent paper clearly showed that hyper-stable “ribbons” were composed of four protofilaments to which MIP4 is bound^33^, suggesting MIP4 stabilizes the ribbon structure. However, no mutant lacking MIPs in A-tubules has been isolated based on the swimming phenotypes of *Chlamydomonas*, implying that MIPs in A-tubules might also have fail-safe mechanisms, similar to MIP3a and c.

## Materials and Methods

### Strains, culture conditions, and isolation of *fap45* and *fap52* mutants

The *Chlamydomonas* strains used in this study are listed in Supplementary Table S3. The double and triple mutants were constructed by standard methods^3^. All cells were cultured on Tris-acetate-phosphate (TAP) plates with 1.5% agar or in TAP medium. *fap45* and *fap52* mutants were isolated from a library of mutants generated by *aphVIII* gene insertion^25^. The mutants were backcrossed with wild type several times before use.

### Motility assay using the CLONA system

To assay the motilities of *Chlamydomonas* cells in TAP medium or TAP with ficoll, videos were recorded using a high-speed camera (EX-F1; Casio) attached on a dark-field light microscope (BX51; Olympus) at 600 fps. The videos were analyzed using the CLONA system^34^ and the results were plotted using R Project software.

### Antibodies

The antibodies used in this study are described in Supplementary Table S4. Anti-FAP45 and anti-FAP52 antibodies were raised against full-length FAP45 and FAP52 proteins, respectively. The full-length cDNA of FAP45 or FAP52 was cloned between the EcoRI and BamHI sites of pMAL-c2x (New England Biolabs). In both constructs, a 6×His tag was inserted into the HindIII site of pMAL-c2x to enable purification in two steps. For affinity purification, each cDNA was also cloned between the NdeI and EcoRI sites of pColdI (Takara). Expression of the recombinant proteins was induced in *Escherichia coli* BL21(DE3) with 0.3 mM IPTG, and almost all of the expressed protein was solubilized from each construct. MBP-FAP45-6×His and MBP-FAP52-6×His were purified with amylose resin (New England Biolabs) and then further purified with Ni-NTA agarose (Qiagen). These purified proteins were used as antigens. Each antibody was affinity purified using polyvinylidene difluoride membranes blotted with FAP45-6×His or FAP52-6×His.

### Chemical crosslinking of axonemes

Isolated axonemes of wild type, *fap45*, and *fap52* were treated with 1-ethyl-3-(3-dimethylaminopropyl)carbodiimide (EDC, Thermo Scientific) in HMEK (30 mM Hepes-KOH, pH 7.4, 5 mM MgSO_4_, 1 mM EGTA, 50 mM K-acetate) for 60 min at room temperature. After quenching the reaction, crosslinked axonemes were analyzed by SDS-PAGE and western blotting with anti-FAP45 and anti-FAP52 antibodies. The crosslinked products were immunoprecipitated with anti-FAP45 antibody following a previously described method^35^. Mass spectrometry analysis (LC tandem MS) of the precipitates was performed at the Proteomics and Mass Spectrometry Facility (University of Massachusetts Medical School).

### Immunofluorescence microscopy

Nucleoflagellar apparatus (NFA) was prepared as previously described^36^. After fixation with 2% formaldehyde for 10 min at room temperature, NFAs were treated with cold methanol (−20°C). The samples were immunostained as previously described^37^. Images were taken with a CCD camera (ORCA-R2; Hamamatsu Photonics) linked to a fluorescence microscope (IX70; Olympus).

### Generation of BCCP-tagged strains

Fragments from the start codon to immediately before the stop codon in the FAP45 and FAP52 genes were amplified by genomic PCR and inserted into pIC2 plasmids^38^. In the FAP52 construct, a 3×HA tag was inserted into the C terminus of FAP52. Each construct was linearized and transformed into *fap45* or *fap52* cells by electroporation.

Streptavidin-Alexa546 staining of the axonemes was performed as previously described.^31^

### Thin-section TEM

Samples for thin-section TEM were prepared as previously described^39^ except that we used 50 mM Na-phosphate, pH 7.0 instead of cacodylate buffer. The samples were observed using a transmission electron microscope (JEM-3100FEF; JEOL) equipped with a 4,096×4,096-pixel CMOS camera (TemCam-F416; TVIPS). All images were taken at 300 keV, with ~3 μm defocus, at a magnification of 40,000 and a pixel size of 2.5 Å.

### Cryo-sample preparation

Purified axonemes in HMDEK buffer (30 mM Hepes-KOH (pH7.4), 5 mM MgSO4, 1 mM dithiothreitol, 1 mM EGTA, 50 mM potassium acetate, protease inhibitor cocktail (Nacalai Tesque)) were incubated with anti-beta-tubulin antibody (T0198; SIGMA) for 15 min on ice, followed by incubation with goat anti-mouse IgG (H+L) 15 nm gold (BB International) and 15 nm colloidal gold conjugated with BSA. The mixtures were loaded onto home-made holey carbon grids and plunged into liquid ethane at −180°C using an automated plunge-freezing device (EM GP; Leica).

### Image acquisition

Grids were transferred into the JEM-3100FEF using a high-tilt liquid nitrogen cryo-transfer holder (914; Gatan Inc.). Tilt series images were recorded at −180°C using a K2 summit direct detector (Gatan) and the serialEM^40^. The angular range was ±60° with 2.0° increments. The total electron dose was limited to 100 e^-^/Å^2^ and the nominal magnification was 6,000x. An in-column energy filter was used with a slit width of 20 eV and a pixel size of 7.1 Å.

### Image processing

Image processing for subtomogram averaging of DMT was carried out as previously described^27, 28^. Tilt series images were aligned and back-projected to reconstruct 3D tomograms using IMOD^41^. Alignment and averaging of the subtomograms were performed using custom Ruby-Helix scripts^42^ and PEET software^7^ to average the 96-nm repeats of DMTs. UCSF Chimera was used for isosurface renderings^43^.

### HS-AFM observation

Flagella were demembranated with 0.5% Nonidet P-40 in HMDEK (30 mM Hepes-KOH, 30 mM Hepes-KOH, pH 7.4, 5 mM MgSO4, 1 mM DTT, 1 mM EGTA, 50 mM K-acetate, protease inhibitor cocktail (Nacalai Tesque)), followed by centrifugation for 2 min. The axoneme pellets were suspended with ATP buffer (HMDEK containing 0.1 mM ATP, 1 mM ADP and 0.5% polyethylene glycol (20,000 mol wt)) and incubated for 1 min at room temperature. Frayed axonemes were then diluted with HMDEK and placed on a mica stage. The AFM images were recorded using a home-built high-speed atomic force microscope^44-46^ at frame rates of 1-2 fps. All observations were performed at room temperature.

## Author contribution

MO and MK designed the experiments. MO performed most of the experiments and analyzed the data. MO and MK wrote the paper. MO and TU performed AFM analyses. HY produced the BCCP-tagged constructs and helped with several EM experiments. TY, HI, and HF provided the *Chlamydomonas* mutant library. KW advised on chemical crosslinking experiments. TA advised on AFM analyses.

## Acknowledgements

We thank Dr. John Leszyk (UMMS) for mass spec analysis and Dr. Brian Dynlacht (NYU) for helpful discussion. MO is a recipient of the fellowship from Japan Society for Promotion of Sciences for young scientists. This work was supported by CREST, the Japan Science and Technology Agency to M.K. and T.A., the Japan Society for the Promotion of Science (JSPS) KAKENHI (Grant number 25120714) and JST ALCA to H. F. The authors have no competing financial interests to declare.

## Movies 1

Swimming of wild type, *fap45, fap52*, and *fap45fap52* in TAP medium.

## Movie 2

Swimming of confluent cultured *fap45fap52* in TAP medium.

## Movie 3

Swimming of wild type and *fap45fap52* in TAP with 7.5% ficoll.

## Movie 4

HS-AFM movie of wild type DMT on a mica surface.

## Movie 5

HS-AFM movie of *fap45fap52* DMT on a mica surface.

## Movie 6

HS-AFM movie of *fap45* DMT on a mica surface.

## Movie 7

HS-AFM movie of *fap52* DMT on a mica surface.

## Movie 8

*fap20*, *fap20fap45*, *fap20fap52*, and *fap20fap45fap52* in TAP.

---

**Table S1:** The *Chlamydomonas* flagellar proteome

| Name | Localization | Total Unique | M+M | Axo | KCl | Reference |
| --- | --- | --- | --- | --- | --- | --- |
| Hydin | Central pair | 69 | 0 | 54 | 6 | <sup>1</sup> |
| CPC1 | Central pair | 56 | 0 | 46 | 10 | <sup>2</sup> |
| FAP42 |  | 54 | 2 | 45 | 6 | Kinase? |
| RIB72 | A-tubule | 44 | 5 | 38 | 8 | <sup>3</sup> |
| FAP46 | Central pair | 38 | 0 | 37 | 0 | <sup>4</sup> |
| PF6 | Central pair | 42 | 0 | 36 | 5 | <sup>5</sup> |
| FAP47 | Central pair? | 35 | 0 | 31 | 1 | <sup>4</sup> |
| FAP50 | A-tubule, N-DRC | 31 | 0 | 31 | 1 | <sup>6</sup> |
| FAP45 |  | 39 | 1 | 30 | 5 | This study |
| RIB43a | A-tubule | 33 | 0 | 29 | 0 | <sup>7</sup> |
| FAP52 |  | 30 | 0 | 28 | 5 | This study |

**Table S2:**
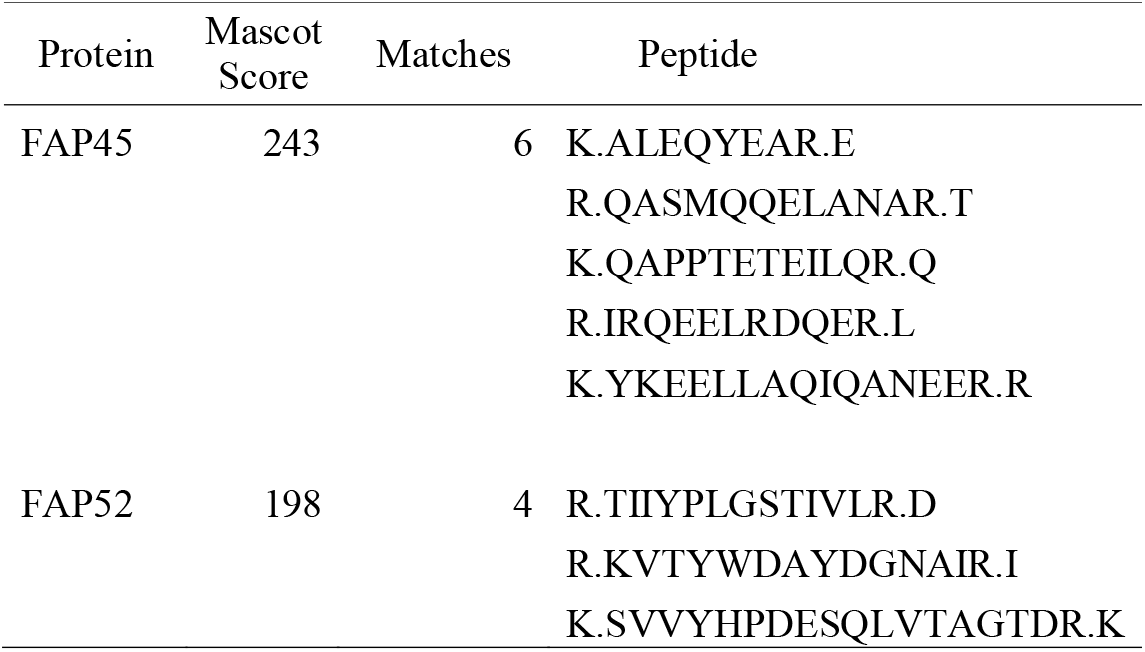
Peptides from the 130 kDa crosslinked product identified by LC MS/MS

**Table S3:** *Chlamydomonas* strains

| Strain | Phenotype | Mutated gene | Reference |
| --- | --- | --- | --- |
| CC124 | Wild type |  |  |
| CC125 | Wild type |  |  |
| *fap20* | Trembling | FAP20 | <sup>8</sup> |
| *fap45* | Slightly slow | FAP45 | This study |
| *fap52* | Wild type | FAP52 | This study |
| *fap45fap52* | slow, paralyzed |  | This study |
| *fap20fap45* | Trembling |  | This study |
| *fap20fap52* | Paralyzed |  | This study |

**Table S4:** Antibodies

| Antibody | Host | Dilution | Reference |
| --- | --- | --- | --- |
| Anti-RSP1 | Rabbit | 1:10000 (WB) | <sup>9</sup> |
| Anti-IC2 (1869A) | Mouse | 1:10000 (WB) | D6168<br>(Sigma-Aldrich) |
| Anti-IC140 | Rabbit | 1:5000 (WB) | <sup>10</sup> |
| Anti-p28 | Rabbit | 1:10000 (WB) | <sup>11</sup> |
| Anti-DRC2 | Rabbit | 1:5000 (WB) | <sup>12</sup> |
| Anti-DRC4 | Rabbit | 1:5000 (WB) | <sup>12</sup> |
| Anti-FAP20 | Rabbit | 1:5000 (WB) | <sup>8</sup> |
| Anti-FAP45 #1 | Rabbit | 1:3000 (WB) | This Study |
| Anti-FAP45 #2 | Guinea pig | 1:200 (IF) | This Study |
| Anti-FAP52 #2 | Rabbit | 1:5000 (WB), 1:200 (IF) | This Study |
| Anti-acetylated α-tubulin (6-11B-1) | Mouse | 1:5000 (WB) | T7451<br>(Sigma-Aldrich) |
| Anti-polyglutamylated tubulin (GT335) | Mouse | 1:10000 (WB) | AG-20B-0020<br>(Adipogen) |

**Supplementary Figure S1.**
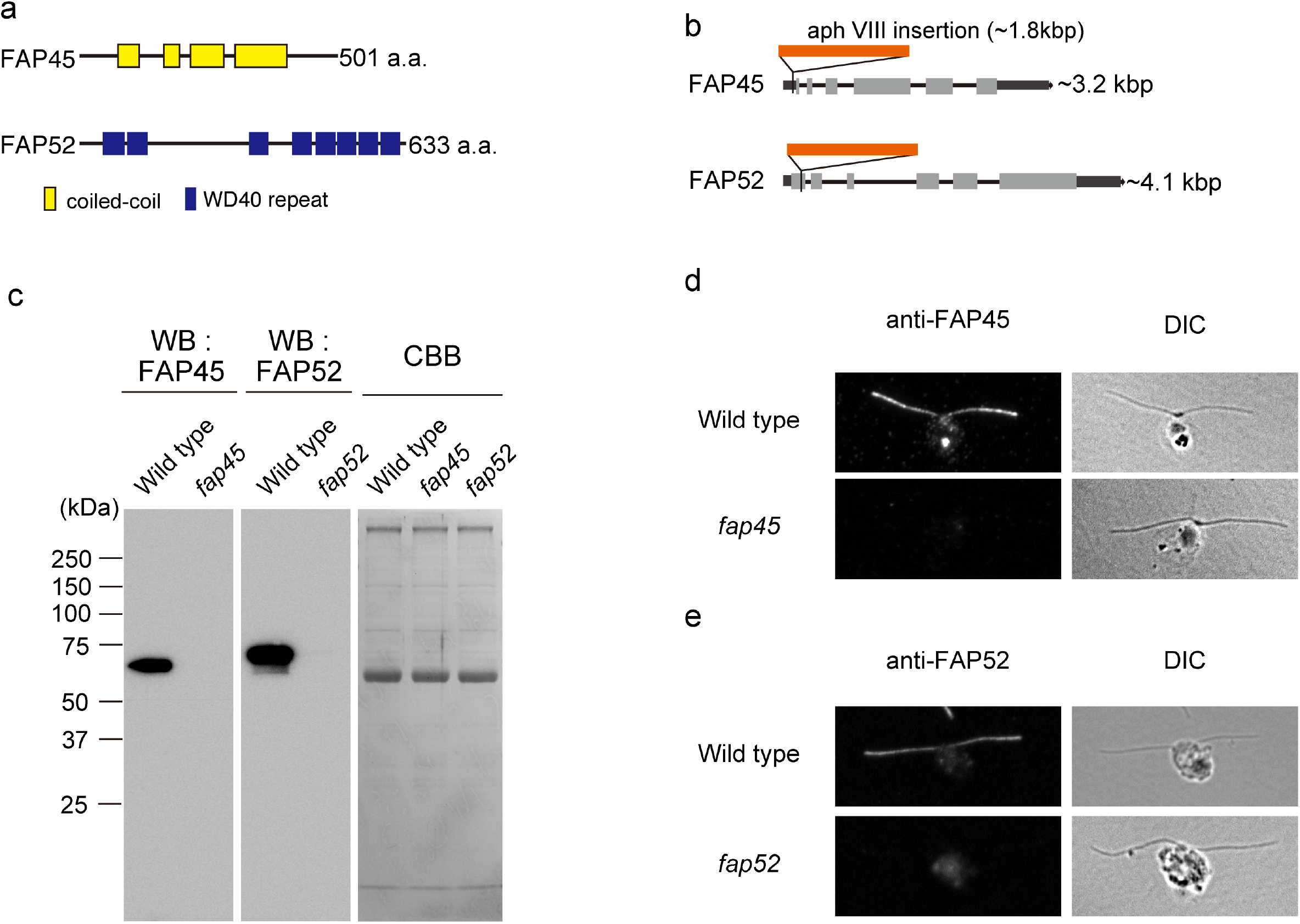
*fap45* and *fap52* are null mutants. (a) Prediction of the structural motifs of FAP45 and FAP52. (b) The insertional mutation sites of *fap45* and *fap52* are indicated on each exon/intron structure. (c) Western blots of axonemes with the anti-FAP45 and anti-FAP52 antibodies. Both antibodies specifically stain the target proteins. (d, e) Immunofluorescence microscopy of the nucleoflagellar apparatus (NFA) stained with the anti-FAP45 antibody (d) and anti-FAP52 antibody (e). Both proteins are distributed along the entire length of wild type axonemes, whereas there is no signal in the mutants.

**Supplementary Figure S2.**
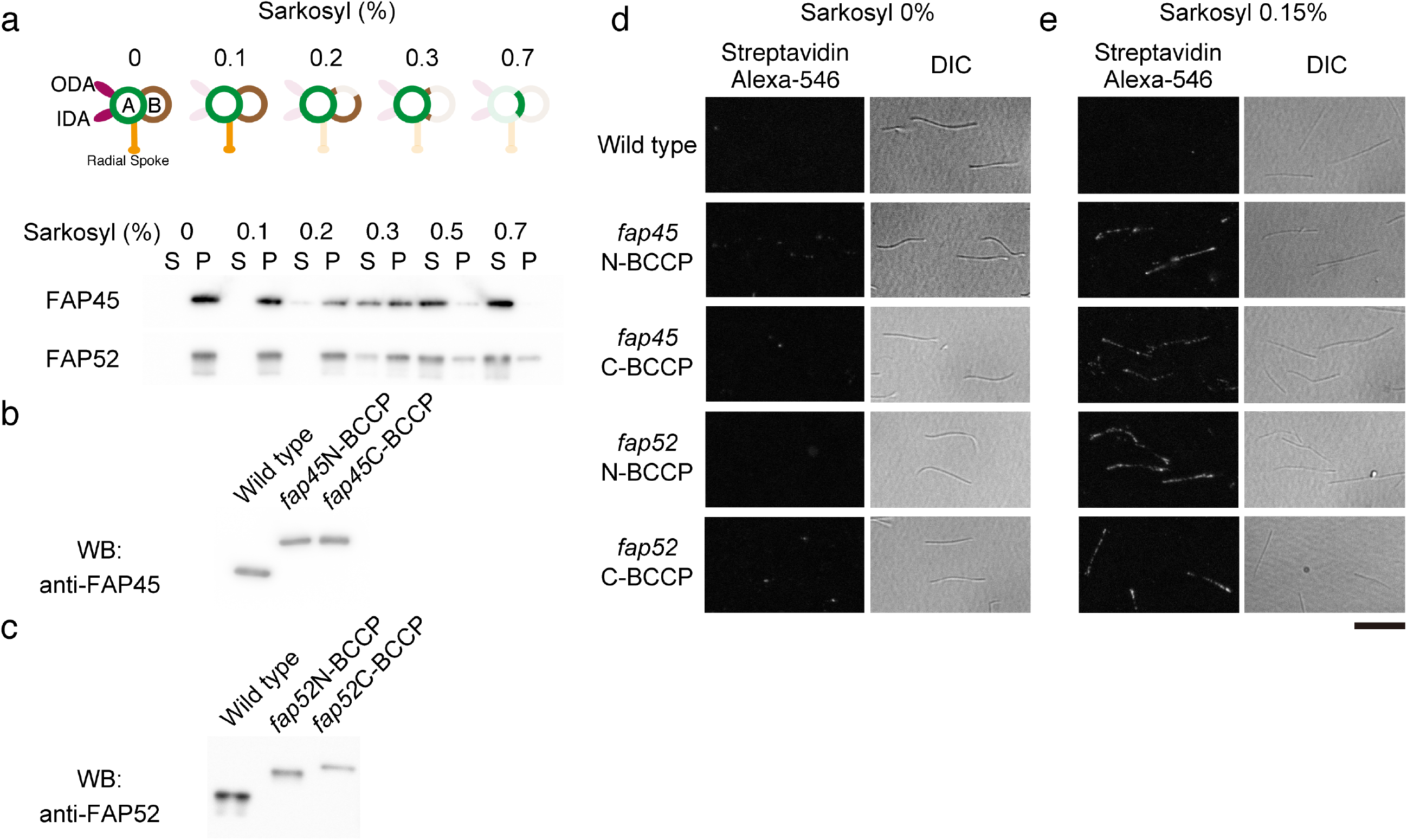
FAP45 and FAP52 are luminal proteins in the B-tubule. (a) Schematic diagrams of the remaining structure of DMT after treatment with various concentrations of sarkosyl. (b) Wild type axonemes were treated with various concentrations of sarkosyl. After centrifugation, the supernatant (S) and precipitate (P) were analyzed by western blot with anti-FAP45 and anti-FAP52 antibodies. (c) Western blot of BCCP tagged FAP45 and FAP52. FAP45 and FAP52 in wild type and BCCP-tagged rescue strains were blotted with anti-FAP45 antibody or anti-FAP52 antibody. (d, e) Axonemes from wild type and BCCP-tagged rescue strains were stained with streptavidin-Alexa546 (d: intact axonemes, e: axonemes treated with 0.15% sarkosyl, DIC: differential interference contrast image, scale bar = 10 μm).

**Supplementary Figure S3.**
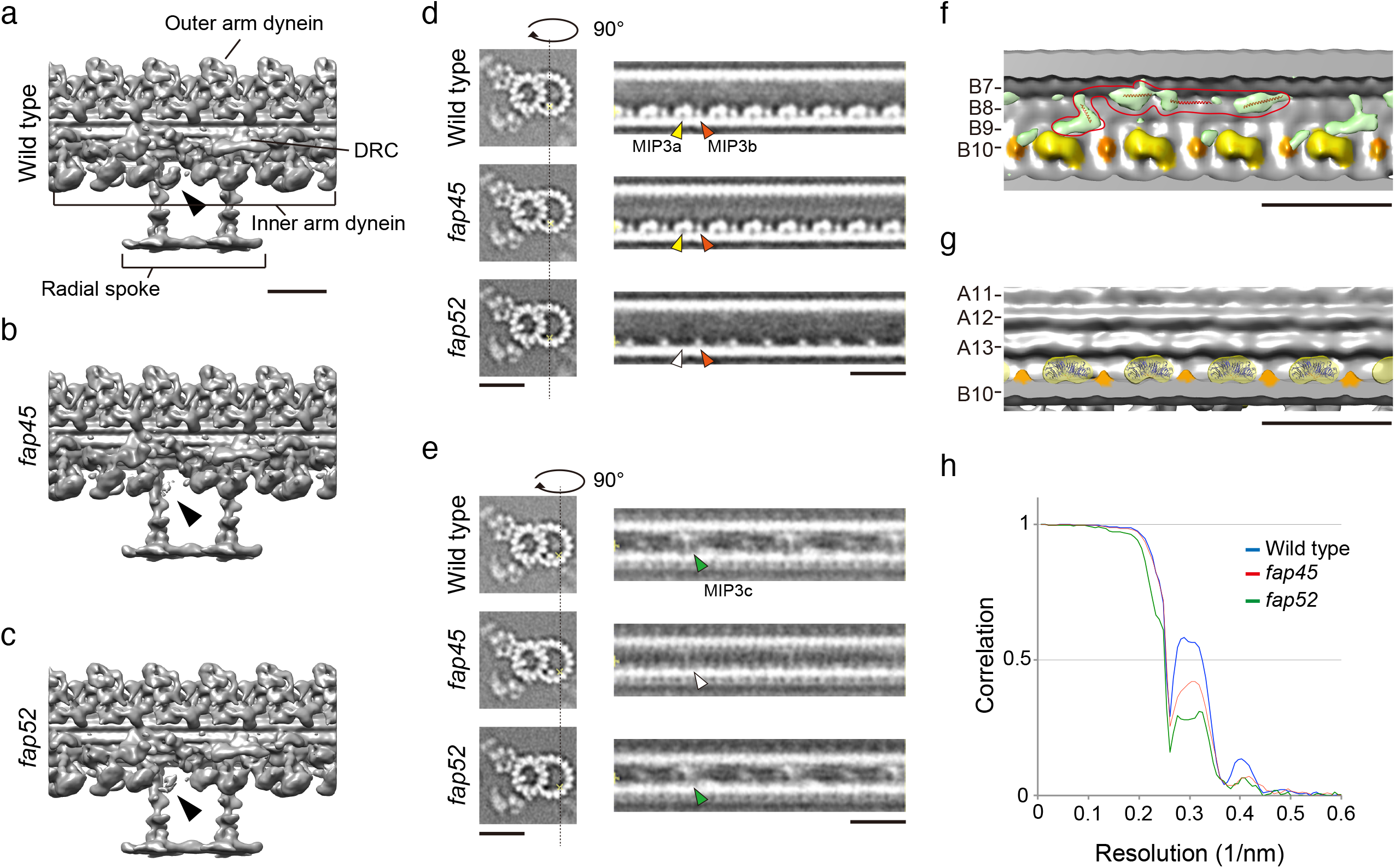
Supporting data of cryo-ET. (a-c) 3D structures of DMTs in wild type (a), *fap45* and (b) *fap52* (c). The density of inner arm dynein e is decreased in *fap45* and *fap52* (arrowheads). Other species of inner arm dyneins, outer arm dyneins, radial spokes, N-DRC are properly arranged on the DMT in *fap45*and *fap52*. (d, e) Density maps of tomographic slices focused on MIP3a and b (d) and MIP3c (e). Scale bar = 25 nm. (f, g) Crystal structures of the coiled coils (PDB: 1DEB, modified) and the WD40-repeats (PDB: 5H1M) were fit to the MIP3c (f) and MIP3a (g) densities, respectively. The *t*-value maps of MIP3a and c were acquired by the Student’s *t*-tests, as previously described^13^‥ The maps circled in red are probable one unit of MIP3c. Scale bar = 25 nm. (h) Fourier shell correlation curves of averaged tomograms used in this study (wild type, *fap45*, and *fap52*). The intersection between each curve and the horizontal line at 0.143 was taken as the effective resolution and is 2.7-2.8 nm.

**Supplementary Figure S4.**
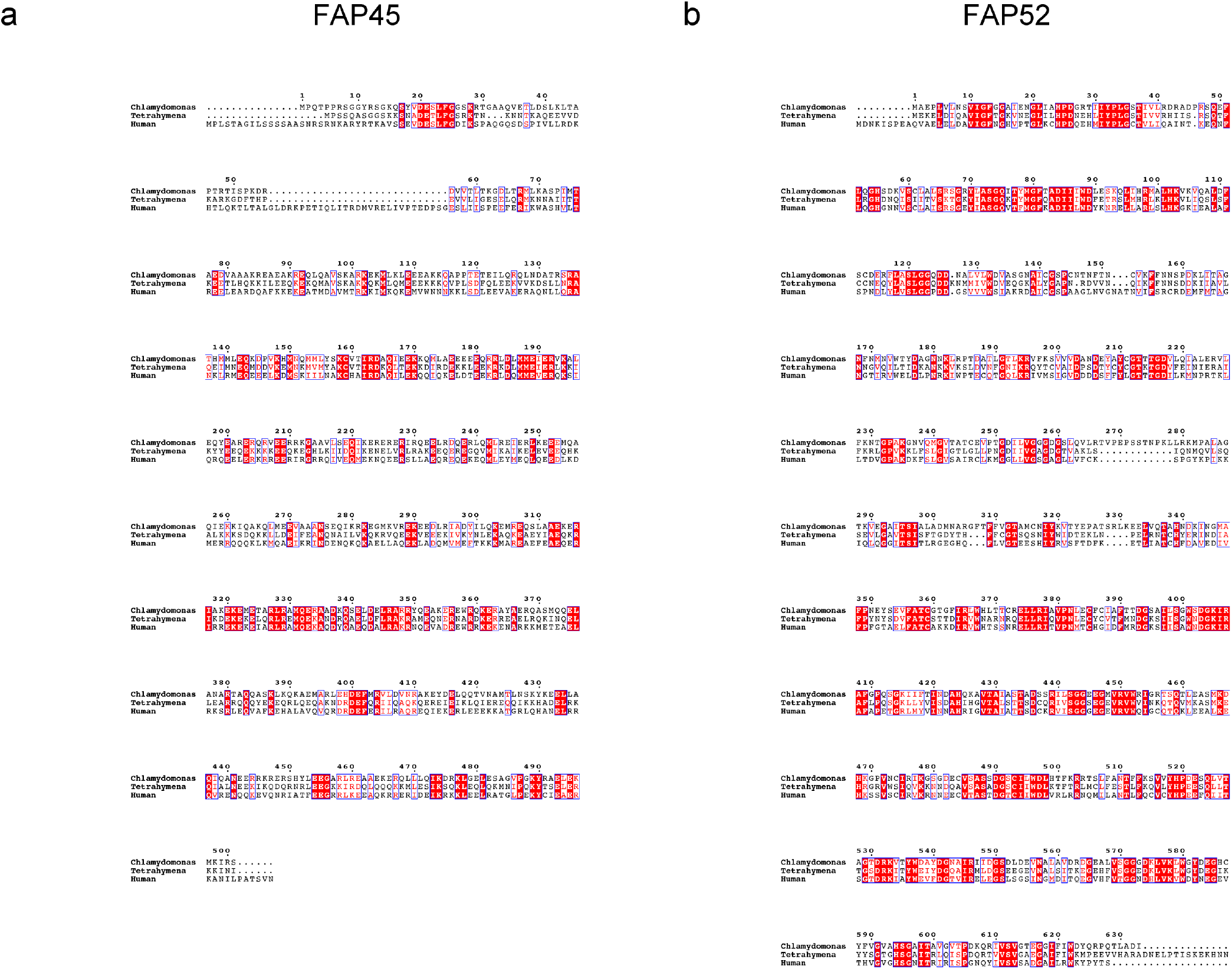
Sequence comparison of FAP45 and FAP52 proteins and their orthologs. *Chlamydomonas* FAP45 (a) and FAP52 (b) were aligned with their homologs in human and *Tetrahymena* using ClustalW (http://www.ddbj.nig.ac.jp/index-j.html). Conserved residues are highlighted with the red box, and similar residues are surrounded by a blue frame. The figures were made using ESPript 3.0 (http://espript.ibcp.fr/ESPript/cgi-bin/ESPript.cgi).

**Supplementary Figure S5.**
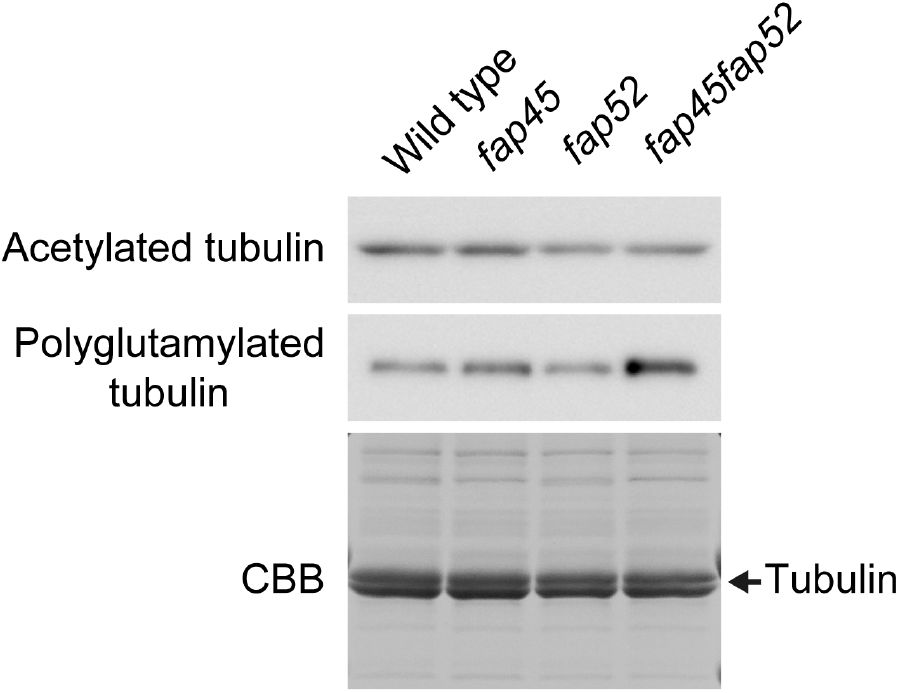
Post-translational modification of tubulin in *fap45*, *fap52*, and *fap45fap52*. Axonemes of wild type, *fap45*, *fap52*, and *fap45fap52* were blotted with anti-acetylated or anti-polyglutamylated tubulin antibody. CBB staining of tubulin is used as a loading control. No significant changes in tubulin acetylation or polyglutamylation were observed in the mutants.

